# Genome-wide DNA polymorphisms in four *Actinidia arguta* genotypes based on whole-genome re-sequencing

**DOI:** 10.1101/694174

**Authors:** Miaomiao Lin, Jinbao Fang, Chungen Hu, Xiujuan Qi, Shihang Sun, Jinyong Chen, Leiming Sun, Yunpeng Zhong

## Abstract

Among the genus *Actinidia*, *Actinidia arguta* possesses the strongest cold resistance and produces fresh fruit with an intense flavor. To investigate genomic variation that may contribute to variation in phenotypic traits, we performed whole-genome re-sequencing of four *A. arguta* genotypes originating from different regions in China and identified the polymorphisms using InDel markers. In total, 4,710,650, 4,787,750, 4,646,026, and 4,590,616 SNPs and 1,481,002, 1,534,198, 1,471,304, and 1,425,393 InDels were detected in the ‘Ruby-3’, ‘Yongfeng male’, ‘Kuilv male’, and ‘Hongbei male’ genomes, respectively, compared with the reference genome sequence of ‘Hongyang’. A subset of 120 InDels were selected for re-sequencing validation. Additionally, genes related to non-synonymous SNPs and InDels in coding domain sequences were screened for functional analysis. The analysis of GO and KEGG showed that genes involved in cellular responses to water deprivation, sucrose transport, decreased oxygen levels and plant hormone signal transduction were significantly enriched in *A. arguta*. The results of this study provide insight into the genomic variation of kiwifruit and can inform future research on molecular breeding to improve cold resistance in kiwifruit.

## Introduction

The genus *Actinidia* includes 52 species and 21 varieties. The *Actinidia chinensis* Planch. species complex consists of large-fruited varieties, such as ‘Hongyang’, that are grown commercially but typically exhibit poor cold resistance. *Actinidia arguta*, which is the second-most widely cultivated *Actinidia* species worldwide, is resistant to cold(A. Ross Ferguson 2007; Latocha and Jankowski 2011; Cossio et al. 2015). *A. arguta* is also the most widespread among all *Actinidia* species and is naturally distributed throughout most of China from the Changbai Mountains in northeast China (latitude 22°N) to the Dawei Mountains in southwest China (latitude 47°N)(Cui 1993; Li et al. 2013). In addition, *A. arguta* exhibits high nutritional value and hairless, edible skin, thus representing an excellent germplasm for breeding improvement. Cold hardiness is a quantitative trait induced by low temperature, and many cold-inducible genes are regulated by *CBF* transcription factors. In a previous study, *CBF* was cloned from the fruit of kiwifruit plants, and the expression of *CBF* was found to be increased when fruit was stored at low temperatures(Ma et al. 2014). Proteomics analysis showed differences in proteins involved in photosynthesis, sugar metabolism, gene regulation, signal transduction, and stress resistance under low-temperature stress in *A. arguta* leaves(Shi et al. 2013). Despite the extensive knowledge regarding the cold-related changes in *A. arguta*, the genetic components underlying these differences remain poorly understood, and associated genomic information for this species is lacking.

The basic chromosome number of kiwifruit (x = 29) is high compared with those of other horticultural crops, and the genus presents extensive inter-taxal and intra-taxal variations in ploidy(A. Ross Ferguson 2007). The sequence of the kiwifruit variety ‘Hongyang’, which was the first species in Ericales to be sequenced, represents a valuable resource not only for biological discovery and crop improvement but also for evolutionary and comparative genomic analysis. The sequence assembly covers ~80% of the estimated genome size of 758 Mb (ME 1994), and its annotation revealed 39,040 predicted genes(Huang et al. 2013). The genome annotation data of ‘Hongyang’ were updated in 2015, including 20 genes that were revised and 30 genes that were created (http://bdg.hfut.edu.cn/kir/index.html). Mining of microRNAs in the ‘Hongyang’ genome and transcriptome has led to the identification of 58 putative microRNAs in kiwifruit(Avsar. B 2015). Li utilized this genome sequence to profile the biosynthesis and accumulation of anthocyanins(Li et al. 2015); however, compared with the genomes of other model plants, the study of the kiwifruit genome is still in its infancy, and little is known about the genomes of other species.

The advent of next-generation sequencing (NGS) technologies has contributed to highly efficient determination of genome-wide genetic variation and genotyping through large-scale re-sequencing of whole genomes. More than 100 plant genomes, ranging in size from 64 Mb to over 5 Gb, have been sequenced to date(Michael and VanBuren 2015). These genomes include those of a number of horticulturally important fruit crops, such as apple(Velasco et al. 2010), grape(Jaillon et al. 2007), Chinese white pear(Wu et al. 2013), papaya(Ming et al. 2008), strawberry(Shulaev et al. 2011), and peach(Cao et al. 2014). In one study, 4.6 million single nucleotide polymorphisms (SNPs) were identified in 74 peach cultivars, including 10 wild varieties, via re-sequencing(Cao et al. 2014). In another study, the genome-wide sequences of two apple cultivars were determined and analyzed to identify floral-associated traits(Xing et al. 2016). Furthermore, numerous SNPs and structural variations (SVs) have been detected in grape through re-sequencing, allowing the discovery of ripening-related genes(Xu et al. 2016). SNPs were first identified in kiwifruit using expressed sequence tag (EST) libraries. The frequency of SNPs in kiwifruit is estimated to be 2,515 SNPs/Mb, and a total of 32,764 SNPs were detected from a combination of four main species and seven different tissues(Crowhurst et al. 2008). A previous study identified a total of 12,586 SNP markers using double-digest RAD sequencing (ddRADseq) (Scaglione et al. 2015). Although the *A. chinensis* genome has already been sequenced and annotated, the absence of diversity within the genome, including a limited number of SNPs and insertions or deletions (InDels), complicates molecular breeding and the identification of target traits. In this study, we re-sequenced four *A. arguta* varieties using the Illumina platform, compared sequence variations with the reference genome of ‘Hongyang’, and analyzed SNPs and InDels. This investigation of whole-genome variations improves our understanding of the cold resistance mechanism of kiwifruit at the molecular level and provides information regarding quantitative trait loci (QTLs) that are associated with cold resistance and can be used in molecular-assisted selection breeding.

## Materials and methods

### Plant Materials

The experimental materials used for re-sequencing included four *A. arguta* genotypes (2n = 4x = 116). The ‘Ruby-3’ (female) genotype and the ‘Hongbei male’ genotype were obtained from Henan Province (113°N, 34°E), China. The ‘Kuilv male’ genotype and ‘Yongfeng male’ were obtained from Jilin Province (125°N, 43°E) and Liaoning Province (123°N, 42°E), China. ‘Hongyang’ (2n = 58) was used as a reference to evaluate cold resistance. All the genotypes were planted at the Zheng Zhou Fruit Research Institute. The leaves of the four *A. arguta* genotypes were collected and used for the DNA extraction and re-sequencing analysis. The shoots of the four *A. arguta* genotypes and the ‘Hongyang’ variety were collected in the dormant period to assess cold hardiness. Eleven genotypes, including 7 *A. arguta* genotypes (‘Ruby-3’, ‘Kuilv’, ‘Xuxiang’, ‘Ruby-4’, ‘Hongbei male’, ‘LD134’, and ‘Hongbei’), 3 *A. chinensis* genotypes (‘Hort16A’, ‘Boshanbiyu’, and ‘Hongyang’), and 1 *A. deliciosa* genotype (‘Hayward’), were used to identify InDels.

### Electrolyte Leakage Tests

One-year-old shoots of the four *A. arguta* genotypes and the *A. chinensis* c.v. ‘Hongyang’ variety were collected in the dormant period, and all the shoots were cut into 20-cm sections and wrapped in plastic film. The samples were placed in a low-temperature incubator (Shanghai Hong Yun Experimental Equipment Factory, Shanghai, China). The *A. arguta* samples were subjected to temperatures of −10°C, −15°C, −20°C, −25°C, and −30°C, whereas the ‘Hongyang’ samples were subjected to temperatures −5°C, −10°C, −15°C, −20°C, and −25°C. The samples were kept at each low temperature for 8 h, followed by a thawing period of 1 h at room temperature. After the freezing treatment, cold hardiness was assessed using the electrolyte leakage method(Zhang and Willison 1987; Odlum and Blake 1996). The lethal temperature at 50% lethality (LT50) was calculated based on the logistic sigmoid function, y = K / (1+ae^−bx^), where y is the REL(Relative Electrolyte Leakage), x is the exposure temperature, a and b are the equation parameters, and k indicates extreme values when x is infinite. The data were calculated and analyzed and standard error were determined using Excel 2013 and SPSS 14.0.

### DNA Library Construction and Sequencing

Young leaf tissues were collected from the four *A. arguta* genotypes for DNA isolation. Total DNA was extracted using the Solarbio DNA Extraction Kit (Beijing Solarbio Science & Technology Co., Ltd, Beijing, China) according to the manufacturer’s instructions. Genomic re-sequencing was performed by Biomarker Technologies (Beijing, China), and the procedure, based on the standard Illumina protocol, was as follows: DNA fragments were generated using ultrasound; the DNA fragments were purified; the ends were repaired with poly-A at the 3′ ends; adaptors were ligated; and clusters were generated. Agarose gel electrophoresis was performed to select specific fragments, and a library was established through PCR amplification. After qualification of the library, sequencing was performed on the Illumina HiSeq 4000 platform.

The raw reads were subsequently evaluated, and low-quality reads (< 20), reads with adaptor sequences, and duplicate reads were filtered. The remaining clean reads were used for mapping. The *A. chinensis* ‘Hongyang’ genome was used as a reference(Huang et al. 2013). The short reads were aligned using the Burrows Wheeler transformation (BWA, 0.7.10 − r789)(Li and Durbin 2009) with the default parameters, except that −M was activated (marking shorter split hits as secondary to make the results compatible with Picard tools software), and the threads for mapping were set to 8 (bwa mem -t 8 -M) to accelerate mapping.

### SNP and InDel Screening

The BWA mapping results were used to detect SNPs and InDels. Picard tools was employed to produce duplicates (http://sourceforge.net/projects/picard/) to limit the influence of PCR duplication. GATK software was used for SNP screening and InDel testing(McKenna et al. 2010). The raw data were translated into sequenced reads through base-calling; during the quality evaluation, the adapters were discarded, and low-quality sequences (quality < 30, or quality by depth < 2.0) were filtered to obtain clean reads. The detected SNPs were screened using the following criteria: coverage depth ≥ 5X, discard the alleles > 2.

### Various Gene Analyses and DNA-Level Functional Annotation

The identified genes with SNPs and InDels were subjected to BLAST searches against functional databases(Altschul et al. 1997). The NCBI non-redundant (NR), Swiss-Prot, Gene Ontology (GO), Clusters of Orthologous Groups of proteins (COG), and Kyoto Encyclopedia of Genes and Genomes (KEGG) annotation databases were used to analyze gene functions(Ashburner et al. 2000; Tatusov et al. 2000; Kanehisa et al. 2004). For the enrichment test, significance was evaluated based on a P-value ≤ 10^−5^ and an FDR value ≤ 0.01.

### InDel Primer Design and Validation

DNA was extracted using the Solarbio DNA Extraction Kit (Beijing Solarbio Science & Technology Co., Ltd, Beijing, China) according to the manufacturer’s instructions. Based on the re-sequencing results, we randomly searched every chromosome for InDels based on an insertion or deletion size ≥10 bp. The InDel primers were designed using Primer Premier 5.0; the parameters for InDel primer design were as follows: PCR product size: 200-500 bp; primer size: 18-24 bp; primer GC content: 40%-60%; and primer Tm: 55-60°C. In total, 120 primers were used, which are listed in Supplementary Table 1. PCR was performed using a PCR mix (Beijing ComWin Biotech Co., Ltd) in a 10-µl reaction volume with the following components: 5 µl of the PCR mix, 0.5 µl of each primer, 1 µl of template DNA, and 3 µl of ddH_2_O. The amplification program consisted of 35 cycles at 94°C for 5 min, 94°C for 30 s, 55°C for 30 s, and 72°C for 30 s, followed by 72°C for 10 min. PCR amplification was performed in a thermocycler (Bio-Rad). The PCR products were tested via 6% polyacrylamide gel electrophoresis (PAGE) under the following conditions: voltage: 80 V, and electrophoresis time: 2 h. Silver staining was performed, and the bands were photographed and analyzed. PopGen32 and NTsys 2.10e software were used to analyze the polymorphisms.

## Results

### Evaluation of Cold Resistance

LT50 is used as a standard index to assess cold hardiness in plants(Ershadi et al. 2015). According to our results, the REL at different temperatures showed an acceptable ‘S’ curve; therefore, LT50 could be calculated using the logistic sigmoid function method. According to the REL curve (Figure 1), *A. arguta* and *A. chinensis* showed an obvious difference at −25°C; the REL was approximately 60% for ‘Hongyang’, and it was approximately 40%-50% for the *A. arguta* genotypes, indicating that the shoots of *A. arguta* are more cold resistant than those of *A. chinensis* ‘Hongyang’ when the temperature is decreased to −25°C. The logistic regression analysis indicated that the LT50 of ‘Hongyang’ was −20.9°C, whereas the LT50 values were −25.0°C, −23.1°C, −31.1°C, and −29.7°C for ‘Ruby-3’, ‘Hongbei male’, ‘Yongfeng male’, and ‘Kuilv male’, respectively (Figure 1). The LT50 values follow the order ‘Yongfeng male’ > ‘Kuilv male’ > ‘Ruby-3’ > ‘Hongbei male’ > ‘Hongyang’. These results indicate that the LT50 value of *A. arguta* is higher in North China than that in Central China, and *A. arguta* exhibits greater cold resistance than *A. chinensis*.

**Figure 1.**
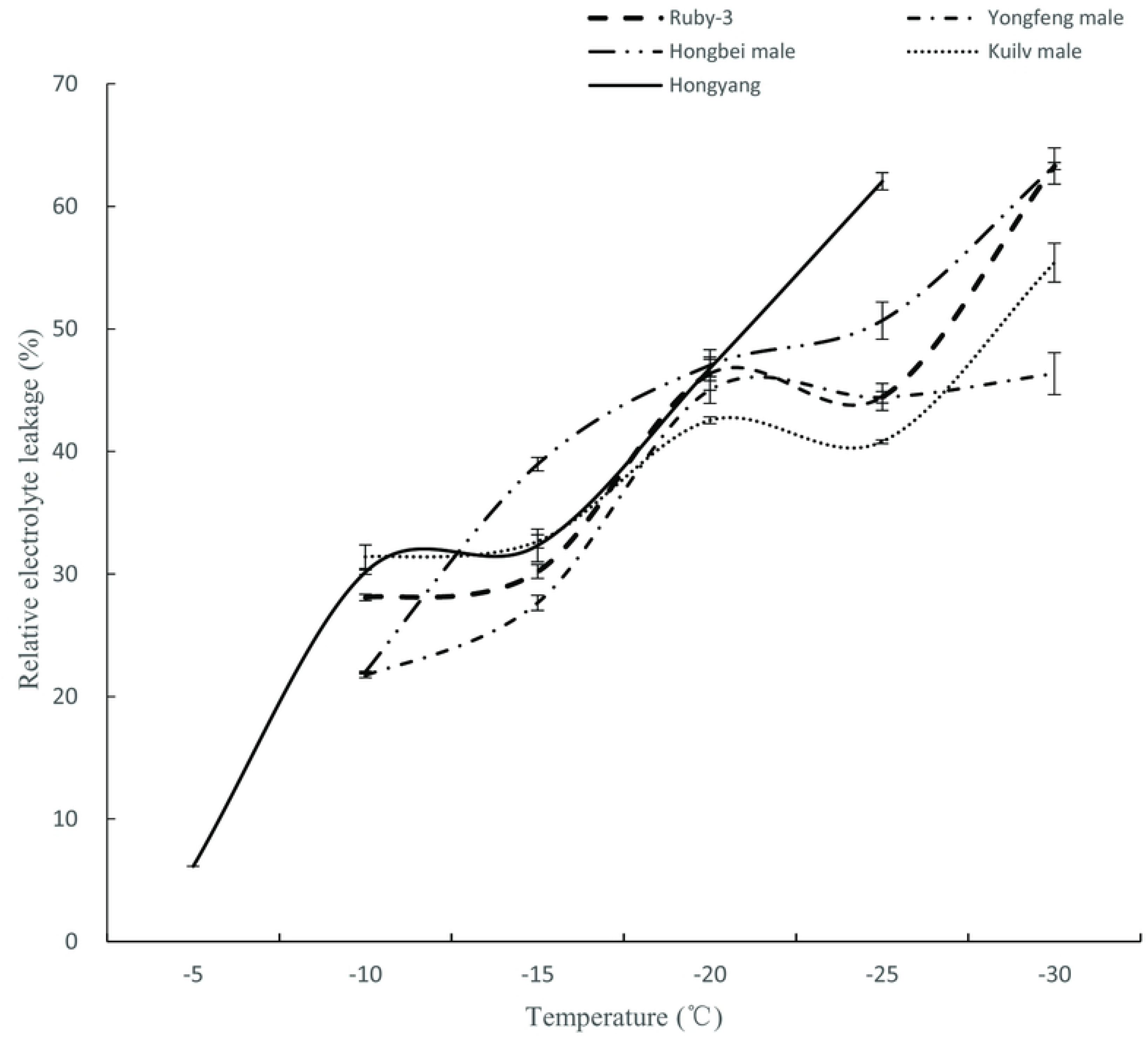
Electrolyte leakage freeze tests were conducted on ‘Hongyang’ and four *A. arguta* genotypes. Data are means (± SE) of three technical replicates.

### Comparison of Reads to the *Actinidia chinensis* ‘Hongyang’ Reference Genome

In this study, we performed genome re-sequencing of four *A. arguta* genotypes as follows. A total of 72,865,383, 58,671,207, 58,281,578, and 69,620,711 clean reads (150 bp) were generated for ‘Ruby-3’, ‘Hongbei male’, ‘Kuilv male’, and ‘Yongfeng male’, respectively; the distribution of each genome was widely uniform, and the sequence was random (Supplementary Figure S1), which indicated good sequence quality. The GC contents were all approximately 38% (i.e., slightly higher than that obtained using the ‘Hongyang’ genome). Approximately 67.68% of the reads mapped to the reference genome (Table 1). The double ‘Hongyang’ genome size was standardized to the *A. arguta* genome size, and the average depths were 16, 14, 15, and 20 in ‘Ruby-3’, ‘Hongbei male’, ‘Kuilv male’, and ‘Yongfeng male’, respectively. All sequencing data for the four *A. arguta* genotypes have been uploaded to NCBI (SRA accession number: SRP118582), and the accession numbers are SRX3209765, SRX3209753, SRX3209750, and SRX3205072 for ‘Ruby-3’, ‘Hongbei male’, ‘Kuilv male’, and ‘Yongfeng male’, respectively.

**Table 1.** Coverage of reads mapped to the reference genome following the re-sequencing of four *A. arguta* genotypes

| Genotype | Raw_Reads | Clean_Reads | GC (%) | Q30 (%) | Mapped (%) | Average_depth |
| --- | --- | --- | --- | --- | --- | --- |
| Ruby-3 | 73,121,307 | 72, 865, 383 | 38.70 | 80.14 | 70.47 | 16 |
| Hongbei male | 58,871,369 | 58, 671, 207 | 38.79 | 82.26 | 68.04 | 14 |
| Kuilv male | 58,498,020 | 58, 281, 578 | 38.33 | 82.73 | 66.43 | 15 |
| Yongfeng male | 69,998,704 | 69, 620, 711 | 38.30 | 80.08 | 65.81 | 20 |

### Analysis of SNPs and InDels

Highly reliable SNPs were identified in the four genotypes. In total, 4,710,650, 4,590,616, 4,646,026, and 4,787,750 SNPs were identified in ‘Ruby-3’, ‘Hongbei male’, ‘Kuilv male’, and ‘Yongfeng male’, respectively. The four genotypes exhibited overlapping and distinct SNPs (Figure 2), with approximately 1 million different SNPs between any two of the genotypes (Table 2), and all the samples displayed different heterozygous and homozygous SNP loci. A greater number of homozygous SNPs corresponds to a greater difference between the samples and the reference genome. The percentages of heterozygous and homozygous SNPs were approximately 30% and 70%, respectively (Table 3). According to the observed nucleotide substitution, the SNP type can be classified as a transition (A/G and T/C) or transversion (A/C, T/G, A/T, and G/C); the transition to transversion ratio (Ti/Tv) in the four varieties was 1.45. The distribution of SNP mutation types showed that C:G > T:A and T:A > C:G, accounting for the high ratio (Figure 3). The distribution of the SNPs in the functional regions of the four varieties was determined and was found to be similar among the varieties. The highest proportion was observed in intergenic regions, which accounted for ~22.63% of the SNPs, followed by downstream regions (15.62%) and upstream regions (13.35%), while SNPs in coding (CDS) regions accounted for 7.95%-8.21% of the SNPs (Table 4). Upstream regions included more SNPs than CDS regions.

**Table 2.**
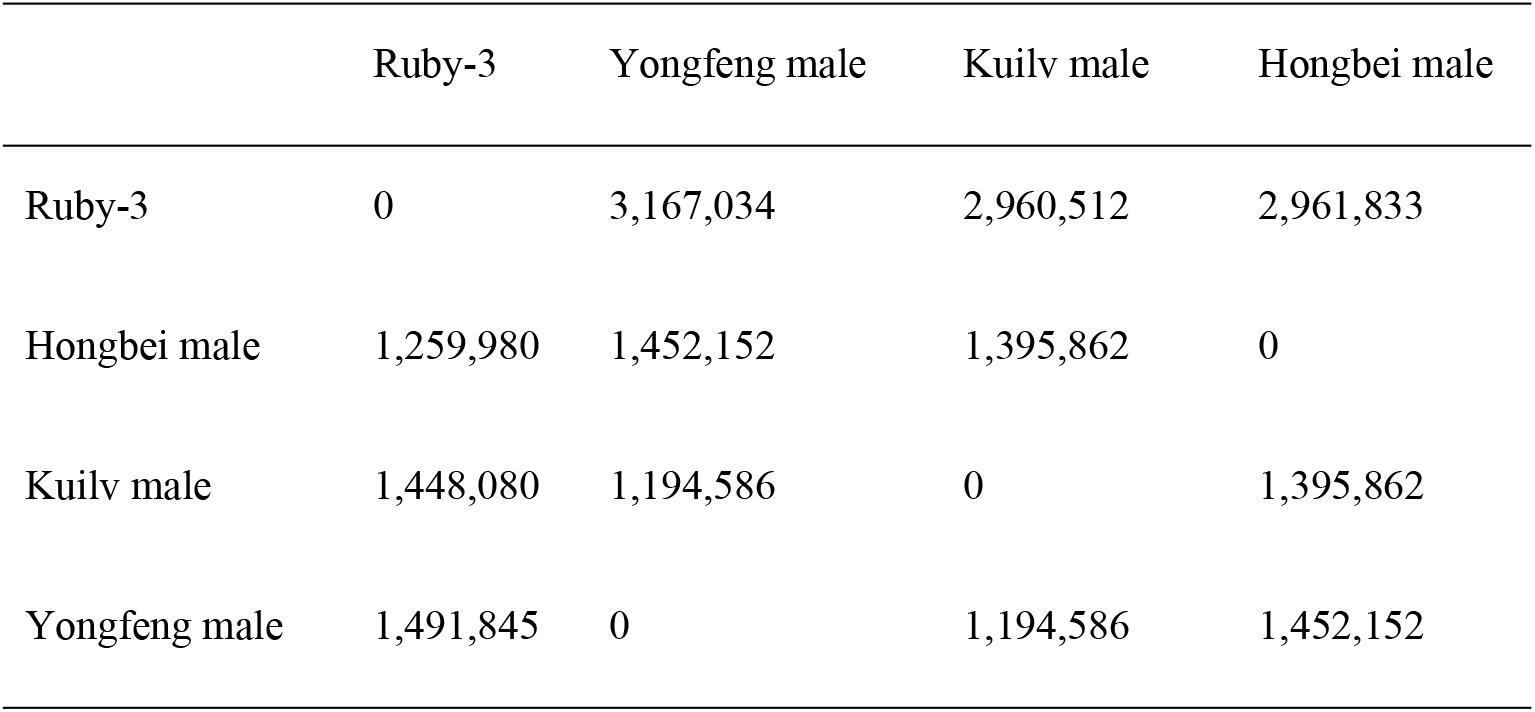
SNP numbers in the four genotypes

**Table 3.** Total number of variants and the type and zygosity of the variants in each genotype

| Genotypes | SNP number | Transition | Transversion | Ti/Tv | Heterozygosity | Homozygosity |
| --- | --- | --- | --- | --- | --- | --- |
| Ruby-3 | 4,710,650 | 2,810,112 | 1,900,538 | 1.47 | 1,345,504<br>(28.56%) | 3,365,146<br>(71.44%) |
| Hongbei male | 3,590,616 | 2,738,635 | 1,851,981 | 1.47 | 1,278,546<br>(27.85%) | 3,312,070<br>(72.15%) |
| Kuilv male | 4,646,026 | 2,769,604 | 1,876,422 | 1.47 | 1,246,789<br>(26.84%) | 3,399,237<br>(75.16%) |
| Yongfeng male | 4,787,750 | 2,855,581 | 1,932,169 | 1.47 | 1,336,943<br>(27.92%) | 3,450,807<br>(72.08%) |

**Table 4.** Ratio of SNP variants in different gene regions of the *A. arguta* genotypes

| Functional regions | Ruby-3 | Hongbei male | Yongfeng male | Kuilv male |
| --- | --- | --- | --- | --- |
| CDS | 15.44 | 15.68 | 15.15 | 15.36 |
| Intergenic | 13.21 | 12.97 | 13.42 | 13.20 |
| Intragenic | 0.00 | 0.00 | 0.00 | 0.00 |
| Upstream | 10.65 | 10.29 | 10.91 | 10.71 |
| Downstream | 11.85 | 11.66 | 12.02 | 11.86 |
| Splice_site_acceptor | 0.02 | 0.02 | 0.02 | 0.02 |
| Splice_site_donor | 0.02 | 0.02 | 0.02 | 0.02 |
| Splice_site_region | 0.74 | 0.75 | 0.72 | 0.73 |

**Figure 2.**
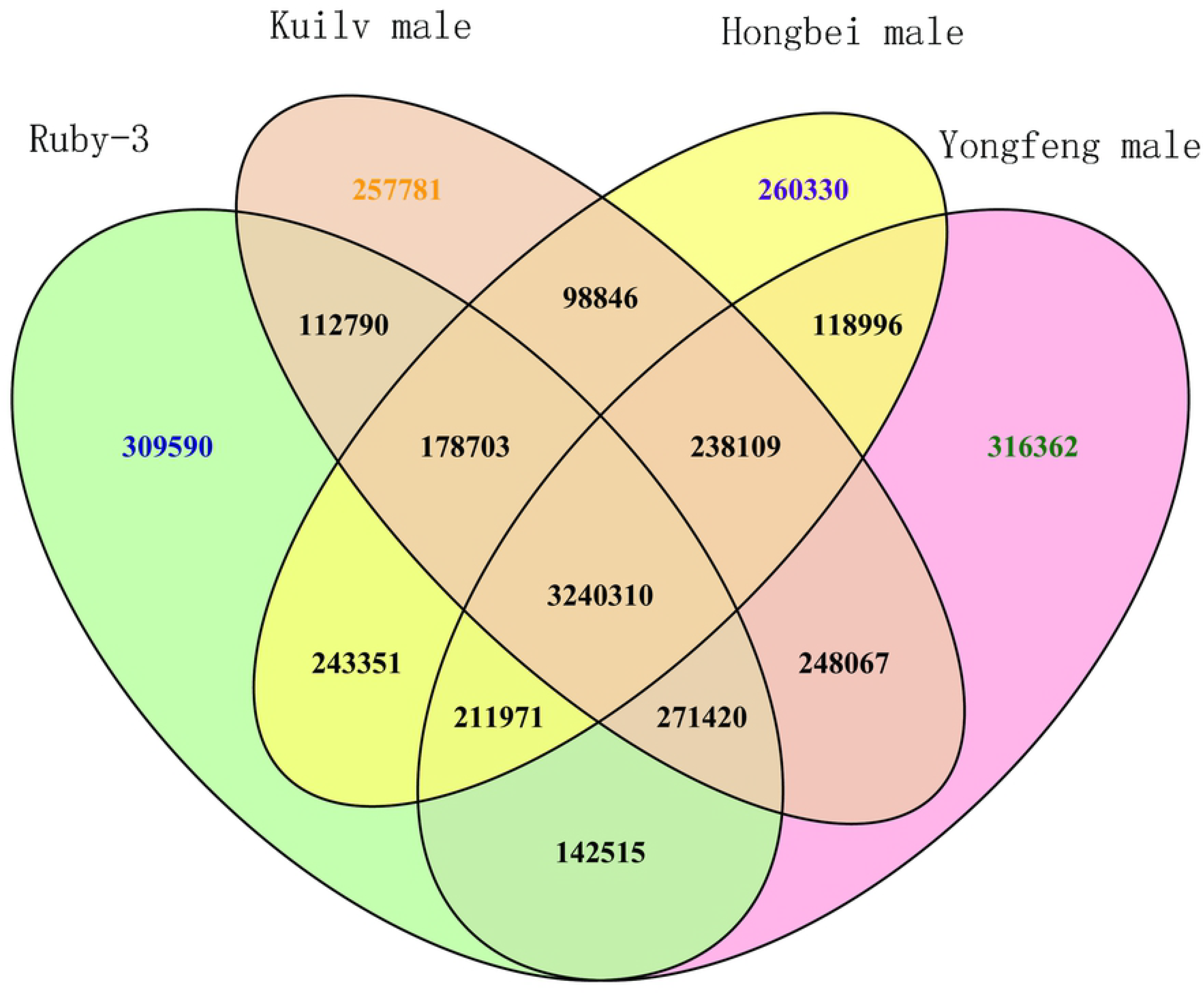
SNP analysis of the four *A. arguta* genotypes.

**Figure 3.**
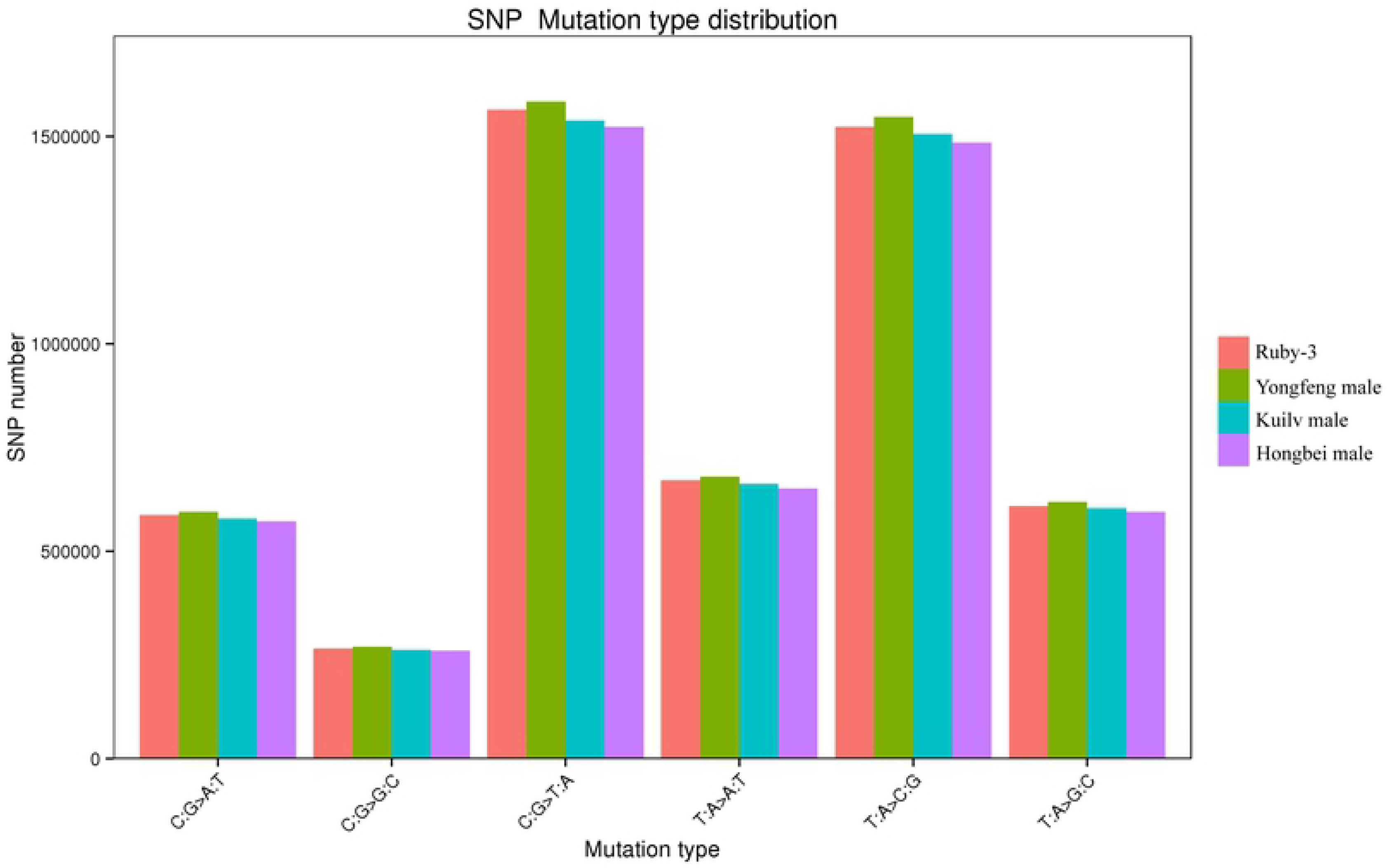
Distribution of SNP mutation types.

In total, 1,481,002, 1,425,393, 1,534,198, and 1,534,198 InDels were detected in ‘Ruby-3’, ‘Hongbei male’, ‘Kuilv male’, and ‘Yongfeng male’, respectively. The types of InDels are presented in Table 5. The InDel length distributions in the whole genome and in the CDS region are shown in Figure 4. Along with InDels longer than 10 bp, InDels with a length of one, two, or three bp accounted for a large proportion of all InDels, accounting for 70.42%, 75.86%, 65.26%, and 70.33% of the InDels in ‘Ruby-3’, ‘Hongbei male’, ‘Kuilv male’, and ‘Yongfeng male’, respectively. A total of 1,005,226 InDels were unique in *A. arguta* relative to ‘Hongyang’ (Figure 5). Approximately 1.75% of the InDels were located in CDS regions. In contrast to the results for SNPs in CDS regions, few InDels were present in these regions, suggesting few insertions or deletions in *A. arguta* relative to *A. chinensis*. However, InDels occupied 15% of upstream regions, including promoter regions (Supplementary Figure 2).

**Table 5.**
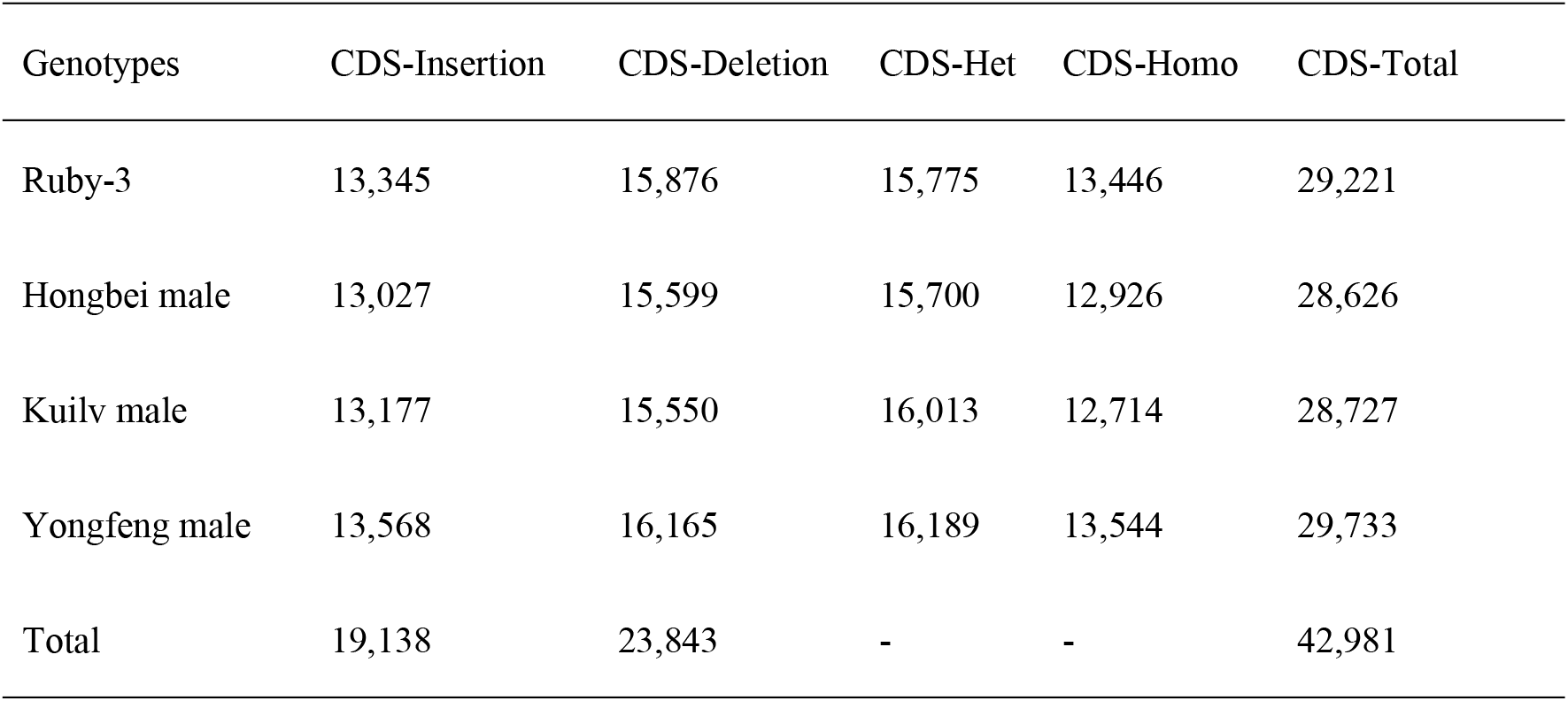
Numbers of InDels of different types in the CDS regions of the four *A. arguta* genotypes

| Genotypes | CDS-Insertion | CDS-Deletion | CDS-Het | CDS-Homo | CDS-Total |
| --- | --- | --- | --- | --- | --- |
| Ruby-3 | 13,345 | 15,876 | 15,775 | 13,446 | 29,221 |
| Hongbei male | 13,027 | 15,599 | 15,700 | 12,926 | 28,626 |
| Kuilv male | 13,177 | 15,550 | 16,013 | 12,714 | 28,727 |
| Yongfeng male | 13,568 | 16,165 | 16,189 | 13,544 | 29,733 |
| Total | 19,138 | 23,843 | - | - | 42,981 |

**Figure 4.**
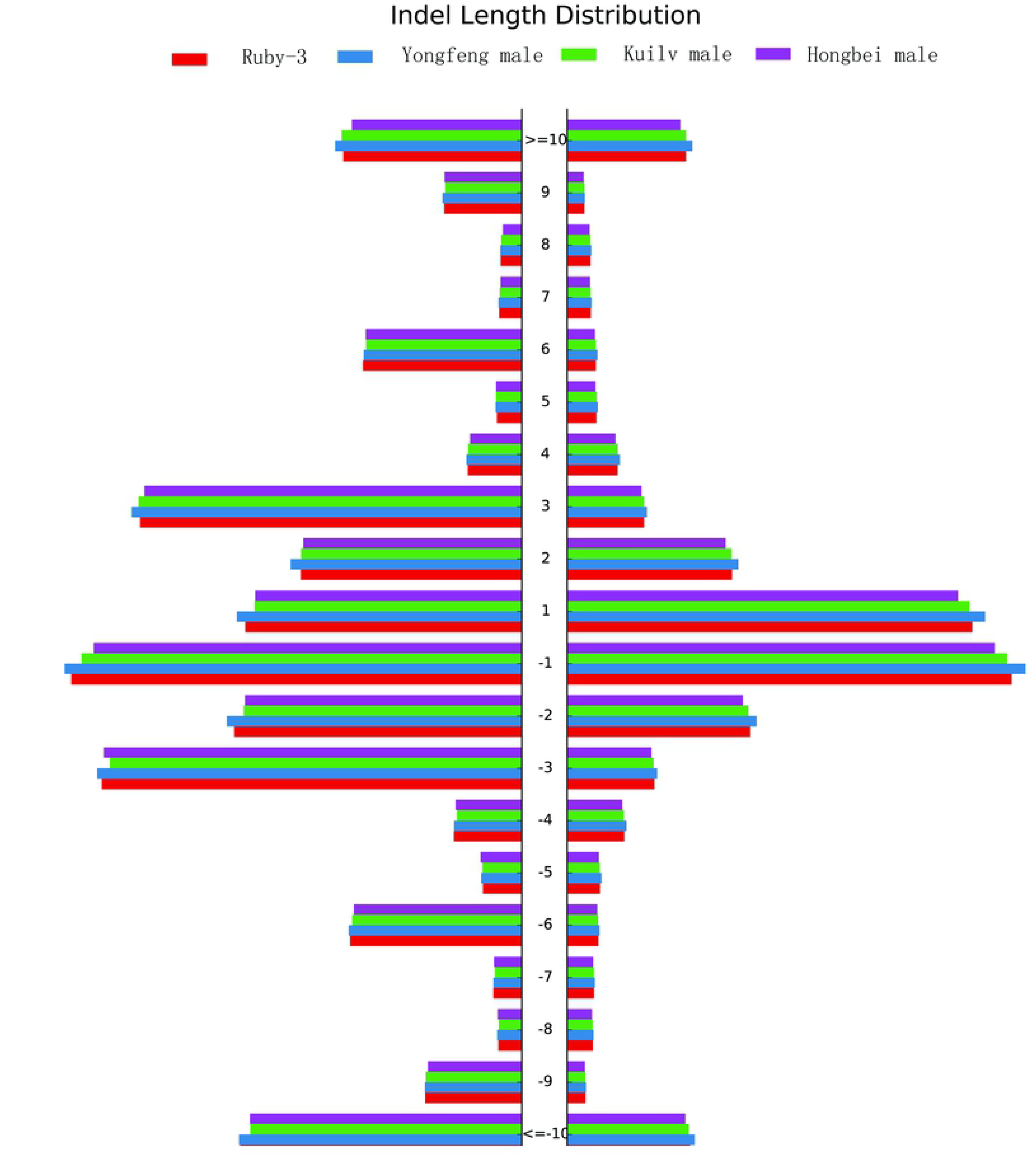
Distribution of different InDel sizes in the four *A. arguta* genotypes.

**Figure 5.**
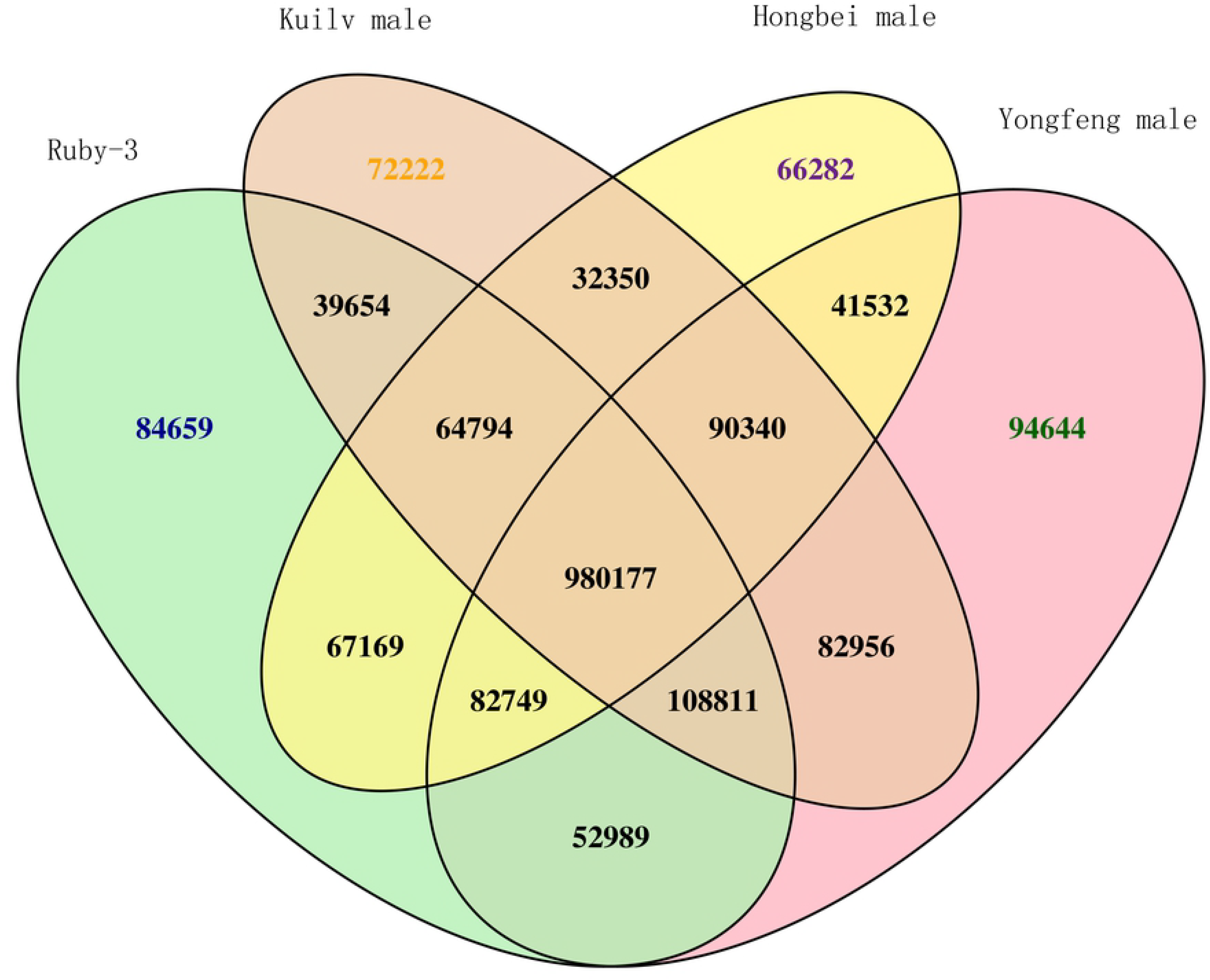
Venn diagram of InDels in the four *A. arguta* genotypes.

### Gene Categories, Functional Annotation, and Differences

Mutations that occur in CDS regions may cause changes in gene function. By examining the non-synonymous SNP mutations and InDels in CDS regions, we identified potential differences in functional genes between *A. arguta* and *A. chinensis*. In total, 22,112, 22,443, 22,077, and 21,935 genes were analyzed in ‘Ruby-3’, ‘Yongfeng male’, ‘Hongbei male’, and ‘Kuilv male’, respectively, using public databases, including the NCBI, NR, Swiss-Prot protein, GO categories, COG, and KEGG databases. Detailed information regarding the functional annotation can be found in Supplemental Table S2. All the functionally annotated genes were classified into GO categories. The GO enrichment classification suggested that the genes from the biological process (BP), cellular component (CC), and molecular function (MF) categories could be divided into 20, 16, and 16 groups, respectively (Figure 6). Based on these categories, a clearer understanding of the genomic characteristics of these kiwifruit genotypes could be obtained. The most abundant components of the BP category were “metabolic process”, “cellular process”, and “biological regulation”. In the CC category, the most abundant components were “cell part” and “cell”, followed by “organelle” and “membrane”. Regarding the BP terms, many genes were classified into the “catalytic activity” and “binding” categories. The GO category analysis also indicated that the genes involved in the cellular response to water deprivation (GO: 0042631), sucrose transport (GO: 0015770), endosome transport via the multivesicular body sorting pathway (GO: 0032509), the response to decreased oxygen levels (GO: 0036293), the response to oxygen levels (GO: 0070482), and the regulation of cellular carbohydrate metabolic processes (GO: 0010675) were significantly enriched in the four genotypes (Supplementary Table S3).

**Figure 6.**
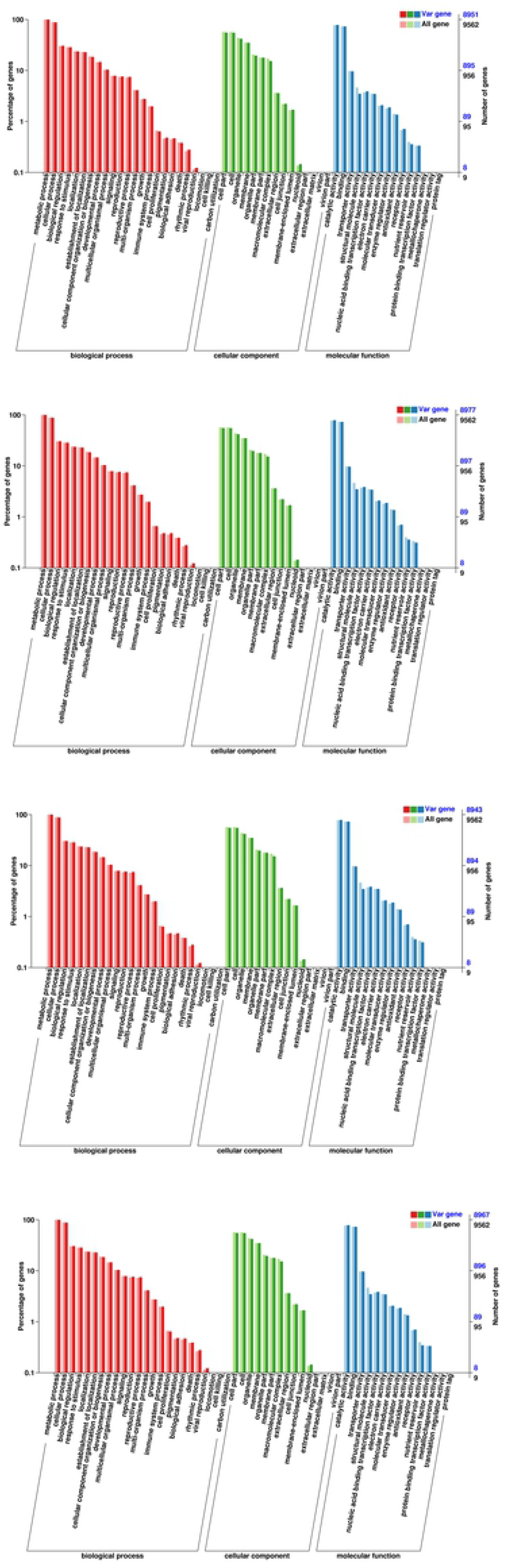
GO classification of differentially expressed unigenes in the four *A. arguta* genotypes. (A) GO classification of differentially expressed genes in ‘Ruby-3’. (B) GO classification of differentially expressed genes in ‘Yongfeng male’. (C) GO classification of differentially expressed genes in ‘Kuilv male’. (D) GO classification of differentially expressed genes in ‘Hongbei male’.

The KEGG pathway analysis showed enrichment of genes involved in 128 pathways, and 13 pathways were significantly enriched (P-value <0.05) (Table 6), including plant hormone signal transduction (ko04075), porphyrin and chlorophyll metabolism (ko00860), and photosynthesis (ko00195). Under the application of a significance threshold of a P-value <0.01, the only metabolic pathway that was enriched was 2-oxocarboxylic acid metabolism. In total, 747, 534, 576, and 674 genes involved in the above pathways were detected, respectively.

**Table 6.** KEGG pathway enrichment analysis

| Pathway | Ko_ID | RB-3 |  | Yongfeng male |  | Kuilv male |  | Hongbei male |  |
| --- | --- | --- | --- | --- | --- | --- | --- | --- | --- |
|  |  | Number of genes | *P*-value | Number of genes | *P*-value | Number of genes | *P*-value | Number of genes | *P*-value |
| 2-Oxocarboxylic acid metabolism | ko01210 | 53 | 0.0067 | 54 | 0.0052 | 51 | 0.0199 | 52 | 0.0137 |
| alpha-Linolenic acid metabolism | ko00592 | 46 | 0.0127 | - | - | 45 | 0.0200 | 44 | 0.0472 |
| Thiamine metabolism | ko00730 | 19 | 0.0138 | 19 | 0.0176 | 19 | 0.0120 | 19 | 0.0140 |
| Fatty acid elongation | ko00062 | 37 | 0.0139 | 38 | 0.0087 | 39 | 0.0017 | 37 | 0.0141 |
| Photosynthesis | ko00195 | 65 | 0.0169 | - | - | 63 | 0.0369 | 64 | 0.0297 |
| Steroid biosynthesis | ko00100 | 27 | 0.0182 | - | - | - | - | - | - |
| Stilbenoid, diarylheptanoid, and<br>gingerol biosynthesis | ko00945 | 26 | 0.0242 | 27 | 0.0123 | 26 | 0.0204 | 26 | 0.0245 |
| Plant hormone signal transduction | ko04075 | 251 | 0.0290 | 265 | 0.0013 | 255 | 0.0048 | 251 | 0.0303 |
| Terpenoid backbone biosynthesis | ko00900 | 54 | 0.0307 | - | - | - | - | 54 | 0.0313 |
| Porphyrin and chlorophyll<br>metabolism | ko00860 | 48 | 0.0312 | 50 | 0.0130 | 48 | 0.0247 | 49 | 0.0168 |
| Zeatin biosynthesis | ko00908 | 31 | 0.0362 | 33 | 0.0092 | 31 | 0.0303 | 31 | 0.0366 |
| Alanine, aspartate, and glutamate<br>metabolism | ko00250 | 47 | 0.0380 | 49 | 0.0161 | - | - | 48 | 0.0209 |
| Flavonoid biosynthesis | ko00941 | 44 | 0.0465 | - | - | - | - | - | - |

### Validation of InDels

To validate the InDels identified in this study, 120 InDels were selected and converted into InDel markers. In the PCR analysis, 81 of the 120 primer pairs exhibited appropriate amplification using genomic DNA from the four *A. arguta* varieties and ‘Hongyang’ as the template; 64 of these 81 primer pairs revealed identifiable polymorphisms among these five varieties based on PAGE analysis. To test the InDel distribution in the other varieties, the diploid *A. chinensis* ‘Boshanbiyu’, tetraploid *A. arguta* ‘Ruby-4’, ‘Hongbei’, and ‘LD134’, and hexaploid *A. deliciosa* ‘Xuxiang’ and ‘Hayward’ genotypes as well as three re-sequenced genotypes (‘Ruby-3’, ‘Kuilv male’, and ‘Hongbei male’) were selected for the identification of InDel polymorphisms (Table 6). In total, 14 InDel markers were selectively amplified (Figure 7), resulting in 43 polymorphic loci among the 11 germplasm resources, with an average of 2.87 loci per primer. Shannon’s diversity index ranged from 0.25 to 0.64, with a mean value of 0.47. The cluster analysis showed that all the varieties could be divided into two categories, with a genetic similarity coefficient of 0.39; one category was composed of *A. arguta*, and the other category consisted of the *A. chinensis* complex.

**Figure 7.**
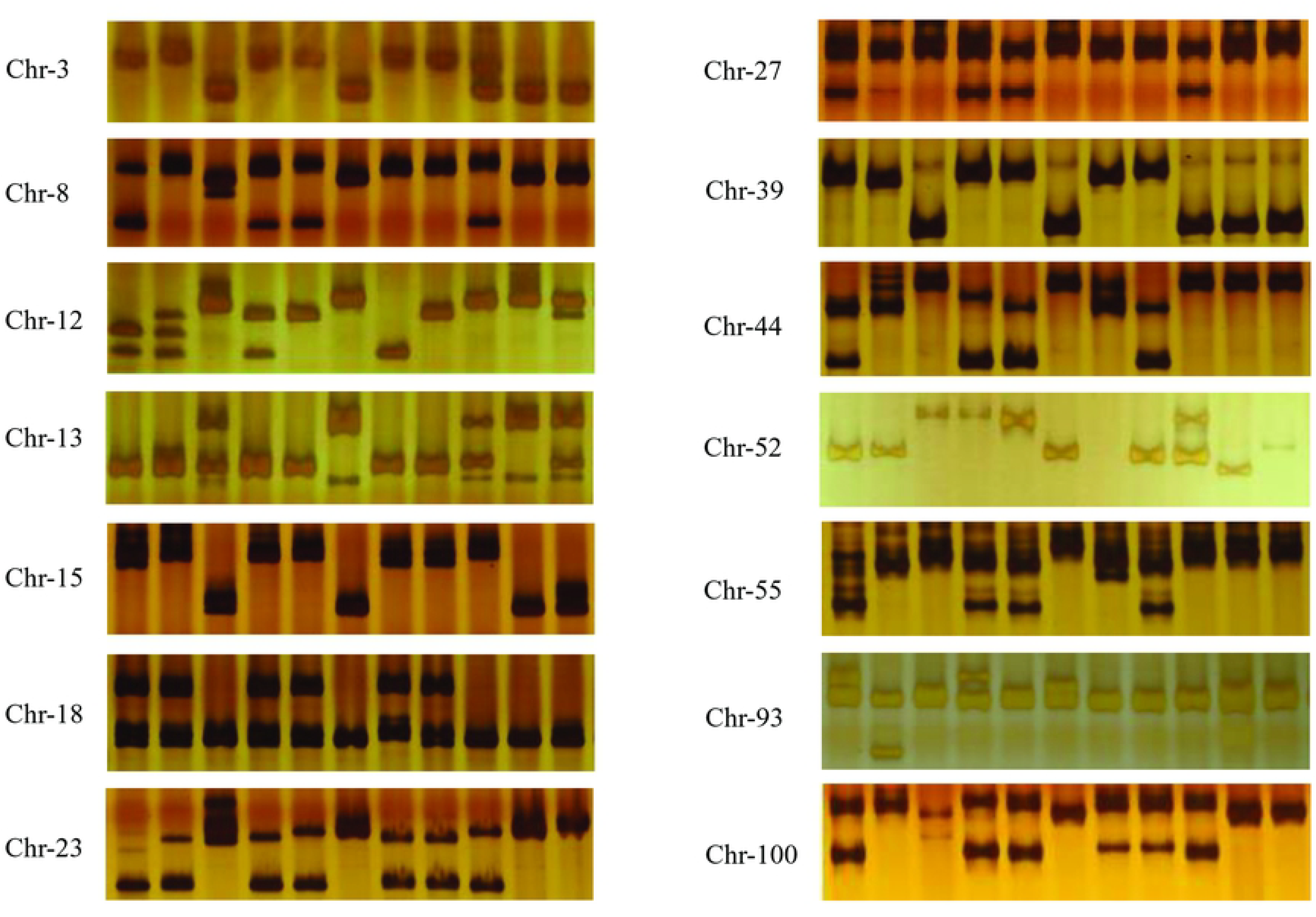
Amplification and electrophoresis segregation of 14 InDel primers in 11 varieties. The varieties from left to right are ‘Ruby-3’, ‘Kuilv’, ‘Xuxiang’, ‘Ruby-4’, ‘Hongbei male’, ‘Hort16A’, ‘LD134’, ‘Hongbei’, ‘Boshanbiyu’, ‘Hongyang’, and ‘Hayward’.

## Discussion

The kiwifruit genome is complex and exhibits variations in ploidy. *Actinidia* species of different ploidies show diverse traits, which are controlled by their unique genomes. The species examined in the present study, *A. arguta* (2n = 4x = 116), also exhibits excellent characteristics. According to our results, the average LT50 of *A. arguta* was −27.2°C, while that of *A. chinensis* ‘Hongyang’ was −20.9°C, indicating that *A. arguta* shows better cold resistance than *A. chinensis*. Chat evaluated plant survival and growth recovery to illustrate that *A. arguta* appeared to be more tolerant to cold than *A. chinensis*(Chat 1995). The examination of cold damage to kiwifruit in the natural environment showed that *A. arguta* exhibited a better survival ability than *A. chinensis* and *A. deliciosa*(Costa et al. 1985), which is consistent with our conclusion in previous work. However, little genetic information is available for *A. arguta*, and this type of information is vital for exploiting molecular markers of desired traits to develop functional genes. In this study, we re-sequenced the genomes of *A. arguta* ‘Ruby-3’, ‘Hongbei male’, ‘Kuilv male’, and ‘Yongfeng male’.

Because genomic information for tetraploid *A. arguta* is lacking, ‘Hongyang’ was used as the reference genome in this study. The mapped reads covered approximately 68% of the reference genome (Table 1); however, whether this variety is autotetraploid or allotetraploid is unclear, and the unmapped sequences may be due to differences in ploidy (e.g., tetraploid *versus* diploid), or the varieties may differ from ‘Hongyang’. Polyploidization events in plants can result in new functions. For example, the sub-genomes of bread wheat display limited gene loss or rearrangement, and cell- and stage-dependent dominance is observed, including in gene families related to baking quality. In *Brassica napus*, however, dynamic shuffling and loss-of-polyploidy events have been reported(Chalhoub et al. 2014), (Pfeifer et al. 2014). In our study, the unknown genome of *A. arguta* is still a large challenge. In this study, SNPs, InDels, and gene functions were analyzed based on the mapped sequence to obtain a better general understanding of *A. arguta*. We analyzed 73% of total reads and found numerous differences in the SNPs and InDels detected in *A. arguta* and *A. chinensis*. Among the SNPs and InDels detected in *A. arguta* and *A. chinensis*, differences were observed in more than 5 million SNPs and 1 million InDels in the different *A. arguta* varieties. These findings offer an overview of the *A. arguta* genome and provide genomic resources for future studies investigating specific characteristics and genetic differentiation. In addition, the results may be particularly useful for the development of excellent cold-resistant trait genes.

The genomic variations identified at the whole-genome level in the four *A. arguta* genotypes that may result in amino acid changes, such as SNPs and InDels, were mainly caused by positive selection during adaptation to environmental changes during the evolutionary process. The observed genome polymorphisms were mainly located in intergenic, upstream, and downstream regions, and similarities were observed among the genotypes (Table 3). Changes in these regions may influence gene expression, but not gene function(Fu et al. 2016). Regarding the substitution of bases, the transition to transversion ratio (Ti/Tv) observed in the four varieties was ~1.45 in our study. This high ratio maintains the structure of the DNA double helix, as shown in rice, in which the Ti/Tv ratio is approximately 2.0-2.5(Wakeley 1996). The ratio of homozygosity to heterozygosity was approximately 2.20 in ‘Ruby-3’ and ‘Hongbei male’ and approximately 2.37 and 2.40 in ‘Kuilv male’ and ‘Yongfeng male’, respectively. *A. arguta* from northern China exhibited greater differences than *A. arguta* from the middle of China. The SNP frequency observed in the current study was about 8,643 SNPs/Mb, which is higher than the reported value of 2,515 SNPs/Mb(Crowhurst et al. 2008), but lower than the value of 15,260 SNPs/Mb observed in *Brassica rapa*(Park et al. 2010). The InDel polymorphism frequency was found to be ~48 InDels/Mb in this study. Among other plants, the frequency of short InDels was found to be 151 InDels/Mb (1-100 bp in size) in *Arabidopsis thaliana* (Jander et al. 2002), approximately 1,050 InDels/Mb in rice(Shen et al. 2004), and 434 InDels/Mb in *B. rapa* when the InDel size was limited to 1-5 bp(Liu et al. 2013), the InDel density was limited to 1-100 bp, and the frequency was limited to 4,830 InDels/Mb (Park et al. 2010). In *A. arguta*, the frequency (1-100 bp) was lower than those observed in these other species. It is likely that the deep sequencing approach used in these previous studies influenced the number of InDels identified.

In the present study, 41 polymorphic loci were detected among the different varieties, and the InDel primers revealed a high rate of polymorphism. The InDel primers were designed based on the re-sequencing data, and 20.8% of the primers were specifically amplified in ‘Hongyang’. In tetraploid and hexaploid species, there are more than two polymorphic loci, which may show different InDels in the sub-genome. The validation data further implied that the InDel primers were effective in different kiwifruit species and may be used to identify phenotypes.

Since the 1980s, *A. arguta* cultivation has been introduced in several regions of Europe, including Belgium, Italy, France, and Iran(Costa et al. 1985; Chat 1995; Van Labeke et al. 2015). In New Zealand*, A. arguta* has shown good cold resistance and is used as a stock for improving cold resistance(Chartier and Blanchet 1997). Qi examined the thickness of the collenchyma and found that collenchyma thickness was closely related to cold resistance(Qi et al. 2011). In the present study, four entire genomes of *A. arguta* were re-sequenced to increase our understanding of plant traits, which could benefit further transcriptome analyses to identify functional genes.

## Acknowledgments

This study was supported by the National Science Foundation of China (Grant No. 31801820) and Special Funds for Science and Technology Innovation Project of Chinese Academy of Agricultural Sciences (Grant No. CAAS - ASTIP - 2018 – ZFRI).

## Author contributions

Conceived and designed the experiments: Jinbao Fang, Chungen Hu, and Miaomiao Lin. Performed the experiments: Miaomiao Lin, Shihang Sun, Yunpeng Zhong, and Leiming Sun. Analyzed the data: Miaomiao Lin, Jinyong Chen and Xiujuan Qi. Wrote the paper: Miaomiao Lin, Chengen Hu, and Jinbao Fang.

